# *<u>De</u> novo* <u>co</u>mpartment <u>de</u>convolution and weight estimation of tumo<u>r</u> samples (DECODER)

**DOI:** 10.1101/561647

**Authors:** Xianlu Laura Peng, Richard A Moffitt, Robert J Torphy, Keith E Volmar, Jen Jen Yeh

## Abstract

Tumors are mixtures of different compartments. While global gene expression analysis profiles the average expression of all compartments in a sample, identifying the specific contribution of each compartment remains a challenge. With the increasing recognition of the importance of non-neoplastic components, the ability to breakdown the gene expression contribution of each is critical. To this end, we developed DECODER, an integrated framework which performs *de novo* deconvolution, and compartment weight estimation for a single sample. We use DECODER to deconvolve 33 TCGA tumor RNA-seq datasets and show that it may be applied to other data types including ATAC-seq. We demonstrate that it can be utilized to reproducibly estimate cellular compartment weights in pancreatic cancer that are clinically meaningful. Application of DECODER across cancer types advances the capability of identifying cellular compartments in an unknown sample and may have implications for identifying the tumor of origin for cancers of unknown primary.

---

Tumor samples are mixtures of distinct cell populations that contribute to intra-tumor heterogeneity, including immune, stroma and normal cells^1,2^. Therefore, with bulk tumor samples, the analysis of tumor gene expression can be significantly confounded by the presence of non-neoplastic cell types, while the contribution of the tumor microenvironment is difficult to separate. Although laser-capture microdissection (LCM) and single cell sequencing techniques strive to tackle these problems, both of them present certain limitations. LCM is labor-intensive and may influence the quality of the microdissected tissue for further analysis^3,4^. Single-cell sequencing is still expensive, computing resource heavy, and currently limited by the lack of comprehensive cell-sorting biomarkers^5,6^.

To eliminate the need of relying on LCM or single-cell-based techniques, a plethora of computational strategies have been developed to deconvolve the mixed signal present in a bulk tumor sample using RNA gene expression, DNA copy number data or DNA methylation data. Algorithms based on DNA copy-number alterations, e.g. ABSOLUTE^7^, and DNA methylation profiles, e.g. MethylPurify^8^ and InfiniumPurify^9^, focus on inferring tumor purity, while expression-based deconvolution methods mainly handle estimation of compartment fractions, as well as extraction of compartment-specific expression profiles^2^. However, the current expression-based deconvolution methods still pose a number of limitations. Some methods are limited to the presupposition of a certain combination of compartments, such as DeMix (tumor and normal)^10^, UNDO (tumor and stroma)^11^ and ESTIMATE (tumor, stroma and immune)^12^. Other methods such as DeconRNAseq^13^ and CIBERSORT^14^, provide the flexibility to measure any number of specific compartments. However, they require knowledge of the pure expression of compartments as the reference, which is difficult to obtain in practice. Similarly, DSA^15^ requires lists of marker genes. However, frequently the exact compartments in a sample are unknown and samples are inherently heterogenous. Therefore, the incomplete knowledge of the number of compartments may hamper the accuracy of calculations of the compartment proportions. In addition, like many other quadratic programming-based algorithms, DSA^15^ has a minimum sample size requirement to perform the estimation, requiring the need for larger datasets.

Pancreatic ductal adenocarcinoma (PDAC) is characterized by relatively low tumor purity and large amounts of desmoplastic stroma. Therefore, identifying tumor-specific alterations in PDAC is a continuing challenge. To perform virtual microdissection and study compartment-specific signatures, we previously deconvolved bulk PDAC samples by adapting the non-negative matrix factorization (NMF) algorithm^16^. As a result, we identified two tumor-specific (Basal-like and Classical) and two stroma-specific (Activated and Normal) subtypes, together with exocrine, endocrine and immune compartments. Like other standard NMF applications, the number of factors (*K*) that the input matrix may be decomposed into must be determined *a priori*. Although the performance of NMF at different *K* may be evaluated by silhouette and cophenetic correlation coefficient, this evaluation assumes the exclusive classification of each sample into one of the *K* clusters^17,18^, which may not be as biologically clear cut. In our previous study, we empirically determined *K* by dedicated manual association of biological relevance to each compartment at multiple trial runs of *K*, which can be time consuming and resource intensive^16^. Thus, developing a streamlined framework that is able to automatically determine *K* is very appealing and will have potential applicability to the bulk tumor sample deconvolution of any cancer type.

Here we present <u>*de*</u> *novo* <u>co</u>mpartment <u>de</u>convolution and weight estimation of tumor samples (DECODER), an NMF-based integrated and automatic framework for the *de novo* deconvolution of tumor mixture samples for compartments, and estimation of compartment weight for samples (Fig. 1a). We show that DECODER can perform automated *de novo* deconvolution for patient samples to derive dynamic and biologically sound compartments without setting ad hoc parameters. By applying DECODER to RNA-sequencing (RNA-seq) data from 33 TCGA (The Cancer Genome Atlas) cancer types, 269 cancer-specific compartments were identified, with a list of marker genes for each compartment. In addition, DECODER can be used to calculate the compartment weights for a single sample, making it potentially applicable in the clinical setting. By applying DECODER to PDAC microarray^16^, TCGA pancreatic adenocarcinoma (PAAD)^19^, COMPASS^20^ and ICGC (International Cancer Genome Consortium) pancreatic cancer^18^ RNA-seq datasets, we demonstrate that DECODER provides insight into pancreatic cancer biology with clinical implications. This framework is not only a useful algorithm for *de novo* and single sample deconvolution of heterogenous samples, but also to the best of our knowledge, for the first time, provides cancer type-specific derived compartment information.

**Fig. 1.**
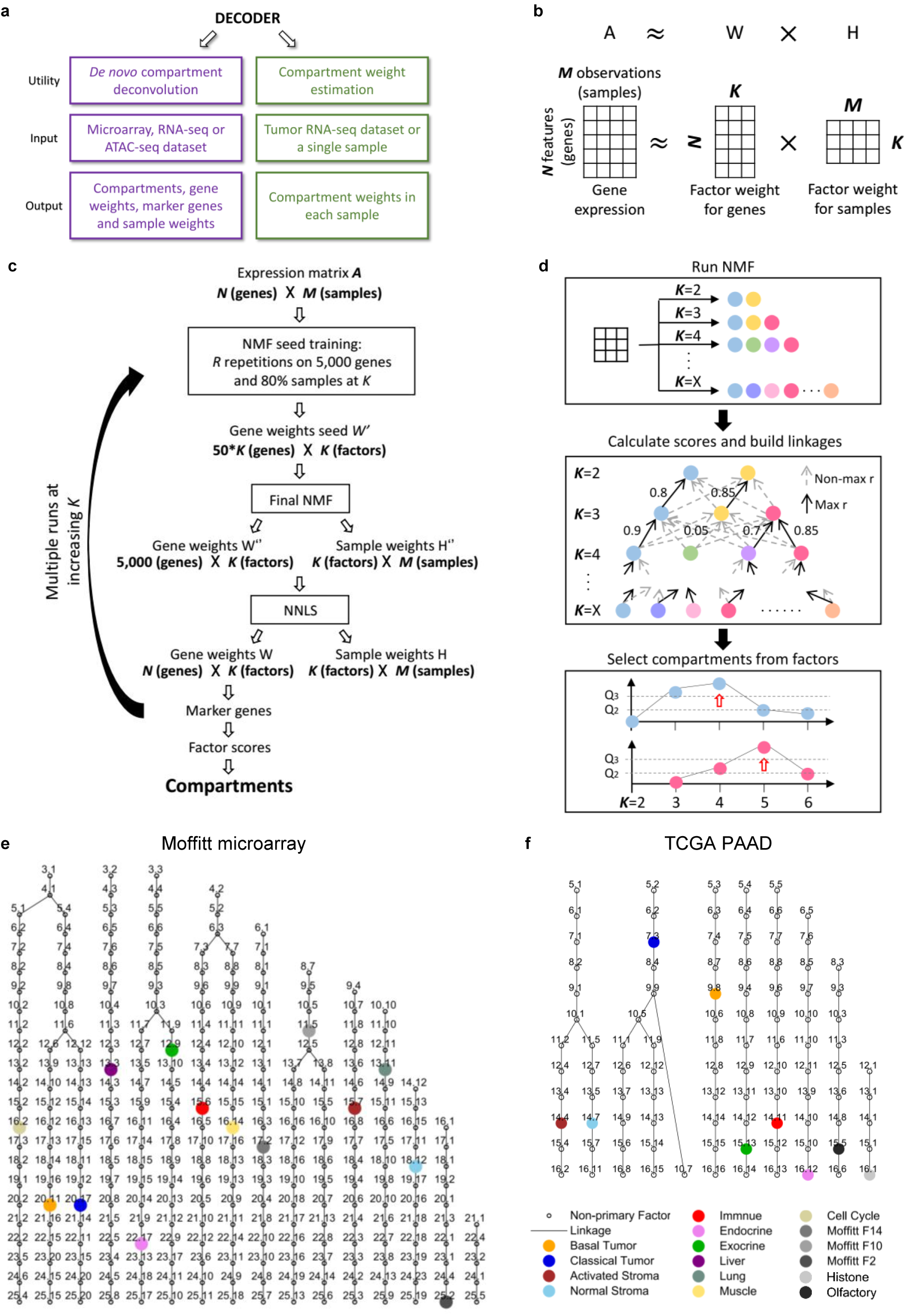
DECODER pipeline. **a** The utility of DECODER includes *de novo* compartment deconvolution for a dataset and single sample compartment weight estimation for TCGA cancers. **b** DECODER uses non-negative matrix factorization (NMF) as the foundational algorithm for de novo deconvolution, where a matrix A is factorized into two matrices W (factor weight for genes or gene weights) and H (factor weight for samples or sample weights). **c** The main steps for *de novo* deconvolution of DECODER. **d** The identification of final compartments from factors in multiple runs of *K*. **e & f** The factor linkage tree for the Moffitt microarray dataset and the TCGA PAAD dataset respectively by applying *de novo* deconvolution of DECODER. The resultant primary compartments were colored, with the dropped factors, secondary and unstable compartments denoted by black circles. Each factor is labeled by the number of factors (K) at current run, a dot and the algorithm assigned random factor number from 1 to *K*.

## Results

### DECODER faithfully captures PDAC compartments in both microarray and RNA-seq datasets

DECODER applies and integrates multiple runs of NMF as the foundational algorithm in the pipeline. Considering a matrix A, consisting of expression levels of *N* genes in *M* samples, the rationale of NMF is to factorize matrix A into two matrices with positive entries, namely A ~ W×H, where W has size *N*×*K* and H has size *K*×*M* (Fig. 1b). *K* denotes the number of factors, with W representing the weight of each gene for each factor (or gene weight) and H representing the weight of each sample for each factor (or sample weight). To avoid *ad hoc* choice of *K* and to automatically determine optimal compartments, DECODER iterates through multiple runs of a carefully designed NMF framework at increasing *K*. In each run, the NMF framework trains a gene weight seed (W’) by *R* iterations of the NMF algorithm followed by applying final NMF and non-negative least squares (NNLS) projection to ensure a robust and reproducible output of W and H at each *K* (Fig. 1c). With the deconvolved factors at multiple runs of increasing *K*, factor linkages are constructed by linking similar factors between adjacent runs based on the percentages of overlapping top genes. Finally, compartments are determined dynamically by evaluating factor scores and score patterns along each branch of linked factors, allowing compartments to be selected at different runs of *K* (Fig. 1d, see Methods).

In a PDAC microarray dataset containing primary tumor, metastatic and normal samples, DECODER identified 14 primary compartments as a blinded determined solution, which accurately reproduced each of the compartments deconvolved previously using NMF at *K*=14 based on empirical trialing^16^ (Fig. 1e). For the matched compartments in the current and previous result, the gene weights show good correlation (median: 0.847, range: 0.761 to 0.924), and the top 250 marker genes show a large overlap of a median of 158 (range: 85 to 207) (Supplementary Fig. S1a,b). The high-level of agreement is reassuring and suggests that DECODER may be used to enable the automated identification of compartments instead of involving labor-intensive manual annotation and empirical determination of *K* at multiple separate runs.

### Deconvolution of 33 TCGA cancer types

We then applied DECODER to TCGA RNA-seq datasets of 33 cancer types separately for *de novo* compartment deconvolution. This *de novo* deconvolution has resulted in the identification of 269 compartments in total, with a median of 9 compartments in each cancer type (range: 4-14) (Supplementary Data 1, complete results available online). Then the ranked list of genes for each compartment was subjected to gene set analysis using the Molecular Signatures Database (MSigDB)^21^ for the biological interpretation and annotation of compartments. The compartments vary from cancer to cancer, while we found five commonly identified ones across cancer types by examining the top gene set terms, namely immune (e.g. IMMUNOLOGICAL_SYNAPSE), basal tumor (e.g. GNF2_SERPINB5), activated stroma (e.g. FARMER_BREAST_CANCER_CLUSTER_5), olfactory (e.g. REACTOME_OLFACTORY_SIGNALING_PATHWAY) and histone (e.g MIPS_UTX_MLL2_3_COMPLEX), which were then manually annotated respectively. Interestingly, even for compartments that were annotated by the same terms, the top genes for different cancer types were found to be different, raising the importance of tumor type specific deconvolution.

For the TCGA PAAD dataset, nine primary compartments were identified. We analyzed each compartment by examining enriched MSigDB gene sets and found that seven dominant compartments in PDAC were reproduced in this dataset, namely basal tumor, classical tumor, activated stroma, normal stroma, immune, endocrine and exocrine, similar to what we found in the microarray dataset (Fig. 1f). For each compartment, by normalizing and ranking gene weights, genes with distinctly large weights were selected as marker genes (Supplementary Fig. S2a, Supplementary Data 2). To investigate the representativeness of the marker genes, we used a linear model described by DSA^15^ to calculate the fractions of seven dominant compartments using these marker genes. We found that the leukocyte fraction and ESTIMATE immune score are correlated with the estimated immune compartment fraction (Supplementary Fig. S2b,c). In addition, both of the ABSOLUTE and methylation based tumor purity were found to highly correlate the sum of the basal and classical compartment fractions (Supplementary Fig. S2d,e), and the ESTIMATE stromal score was found to highly correlate the sum of the activated and normal stroma fractions (Supplementary Fig. S2f). These demonstrated that DECODER can automatically and robustly determine biological compartments in a given dataset *de novo*, with the identification of representative marker genes for each compartment.

### Sample weights (H) derived from *de novo* deconvolution

A matrix of sample weights (H) for compartments was also derived from the deconvolution (Supplementary Data 3). We hypothesized that for each sample, a larger weight is associated with a larger representation for a specific compartment. Indeed, in TCGA PAAD, we demonstrated that normalized DECODER weight for the immune compartment is highly correlated with leukocyte fraction (Fig. 2a) and ESTIMATE immune score (Fig. 2b). Tumor hematoxylin and eosin (H&E) slides were also annotated for the presence or absence of immune infiltrate and tertiary lymphoid structures within the tumor. Samples with high immune weights show apparent infiltration, which is absent in low immune weighted samples (Fig. 2d). Quantitatively, samples containing tertiary lymphoid structures show significantly higher immune weights than those with no tertiary lymphoid structures (Fig. 2c). For tumor compartments, we demonstrated high correlation between the sum of basal and classical tumor weights, and tumor purity based on both ABSOLUTE and methylation (Fig. 2e,f). Similarly, the ESTIMATE stromal score is mirrored by the sum of activated and normal stroma weights (Fig. 2g).

**Fig. 2.**
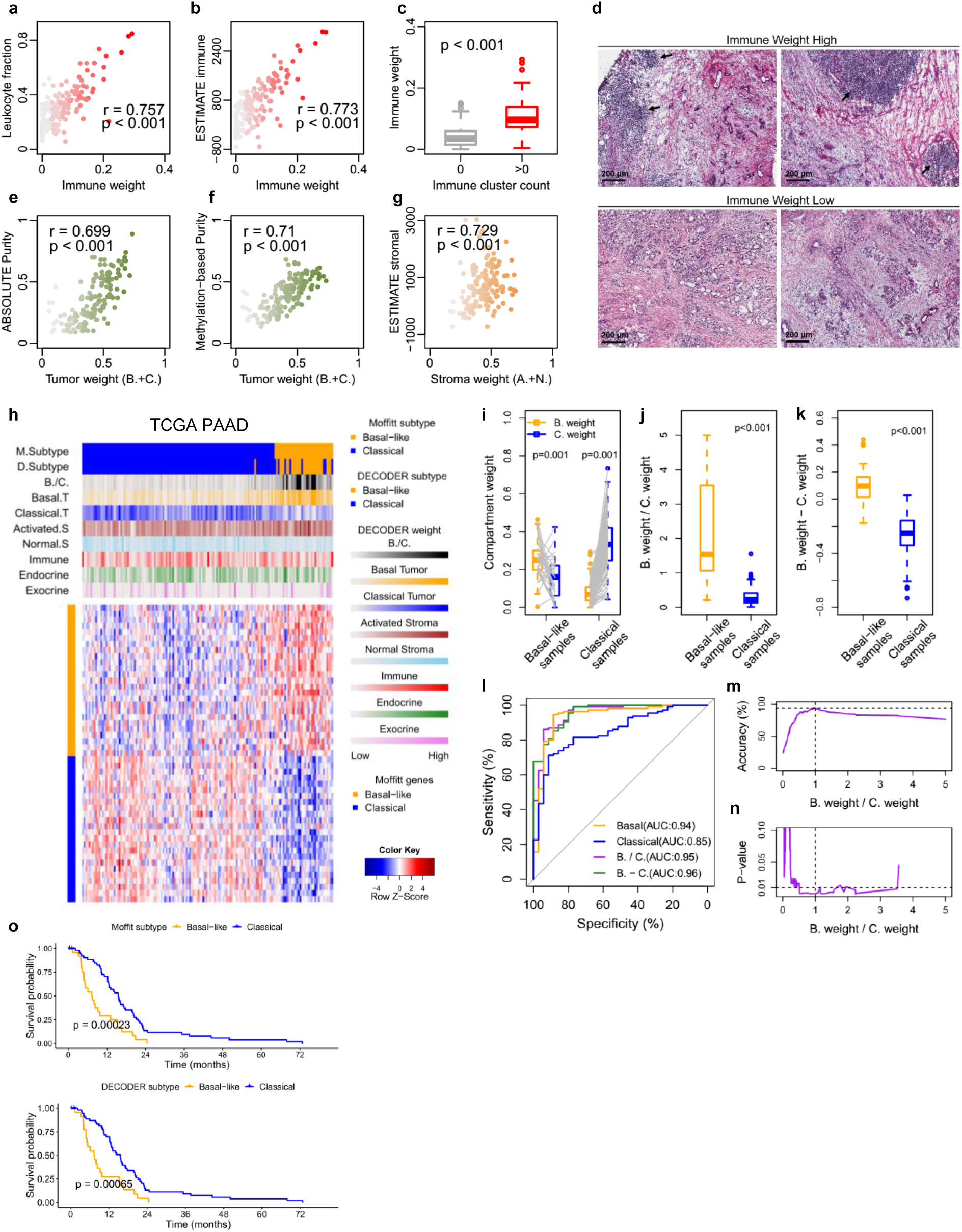
Deconvolved compartment weights for samples in TCGA PAAD samples. **a & b** DECODER immune weight correlations with the leukocyte fraction and ESTIMATE immune score. **c** Quantification of immune clusters by hematoxylin and eosin (H&E) staining were correlated with immune weights. **d** Representative H&E stained PDAC tumor sections showing immune cell infiltration (arrows) in samples with high immune vs low immune weight samples. **e & f** The sum of basal and classical weights correlations with tumor purity estimated by ABSOLUTE and methylation. **g** The sum of activated and normal stroma correlation with ESTIMATE stromal score. **h** DECODER compartment weights, the ratio between basal and classical compartment weights (B./C.), tumor subtypes called by B./C. (D.Subtype) and tumor subtypes called by using Moffitt tumor genes (M.Subtype) are shown for TCGA PAAD samples. Samples are ordered by consensus clustering using Moffitt Basal-like and Classical genes. **i, j & k** Basal weights, classical weights, the ratio (B./C.) and difference (B.-C.) between basal and classical weights in previously called Basal-like and Classical samples. **l** The receiver operating curve (ROC) for basal weight, classical weight, B./C. and B.-C. in classifying subtypes. The area under the ROC (AUC) is shown for each parameter. **m & n** Threshold determination for B./C. to classify subtypes based on accuracy and significance to differentiate survival after classification. **o** Kaplan-Meier plots of overall survival in patients with resected PDAC for Moffitt and DECODER tumor subtypes.

Based on previous consensus clustering based subtype calls for these samples^19^, we show that Basal-like samples are associated with higher basal compartment weights and Classical samples with higher classical compartment weights (Fig. 2h, 2i). Hereafter, for clarity, basal and classical refer to DECODER derived compartments, and <u>B</u>asal-like and <u>C</u>lassical (uppercase) refer to subtypes. Similarly, higher ratios or differences of basal vs classical compartment weights are associated with previous Basal-like subtype calls (Fig. 2j, 2k). To determine the clinical significance of basal vs classical compartment weights, we compared the utility of compartment weights as a clinical variable. We found that either the ratio (*B./C.*) (p=0.002) or difference (*B.-C*.) (p=0.015) between basal and classical compartment weight is associated with survival.

To determine the accuracy of DECODER based subtype calls from basal and classical compartment weights, we calculated the area under the receiver operating curve (AUC) using previous subtype calls as the gold standard. We found that basal compartment weight alone, *B./C.*, and *B.-C*., have similarly high area under the receiver operating curve (AUC 0.94-0.96, Fig 2l). Because *B./C.* shows the second highest AUC after *B.-C*. and is better associated with outcome than *B.-C*., we then determined a threshold for the ratio (*B./C.* = 1) to reach optimal accuracy to call subtypes (Fig. 2m), and optimal significance to differentiate patient outcome (Fig 2n), similar with previous subtype calls (Fig. 2o).

### Single sample weight estimation

Since the sample weights for TCGA dataset were obtained through *de novo* deconvolution which requires a dataset with multiple samples and relatively large amounts of computing time during the training process, we next developed a method to deconvolve RNA-seq samples and calculate the sample weights without the need to apply the *de novo* deconvolution. This method was built using the deconvolved gene weights for compartments and applying NNLS to indicate the sample weights for even a single sample (see Methods). Ten-fold cross validation demonstrated good reproducibility of sample weights derived from gene weights compared to those derived from *de novo* deconvolution (Supplementary Fig. S3a,b). We applied this algorithm to calculate the sample weights for COMPASS and ICGC pancreatic cancer datasets based on gene weights derived in *de novo* deconvolution of the TCGA PAAD dataset (Supplementary Data 3).

Interestingly, in COMPASS, where samples were microdissected, the endocrine, exocrine and immune weights are significantly lower than those in the TCGA PAAD and ICGC datasets (Fig. 3a). In the ICGC dataset, similar to our TCGA PAAD results, we find that ABSOLUTE-based tumor purity, ESTIMATE-based immune and stromal scores correlate with respective DECODER weights (Fig. 3b-d). For the two acinar cell carcinoma samples in this dataset, we find that the exocrine scores are 4.78- and 6.96-fold higher than the mean of the exocrine scores in all other samples. This suggests that the calculated DECODER weights can accurately capture the biological composition of a sample. In addition, in the ICGC dataset, the ADEX subtype samples are associated with higher exocrine weights and the Immunogenic subtype samples are associated with higher immune weights (Fig. 3e). These results may suggest that ADEX and Immunogenic samples are characterized by an overrepresentation of exocrine and immune cells in the respective samples, which may have masked the identification of tumor-specific subtypes^19,22^.

**Fig. 3.**
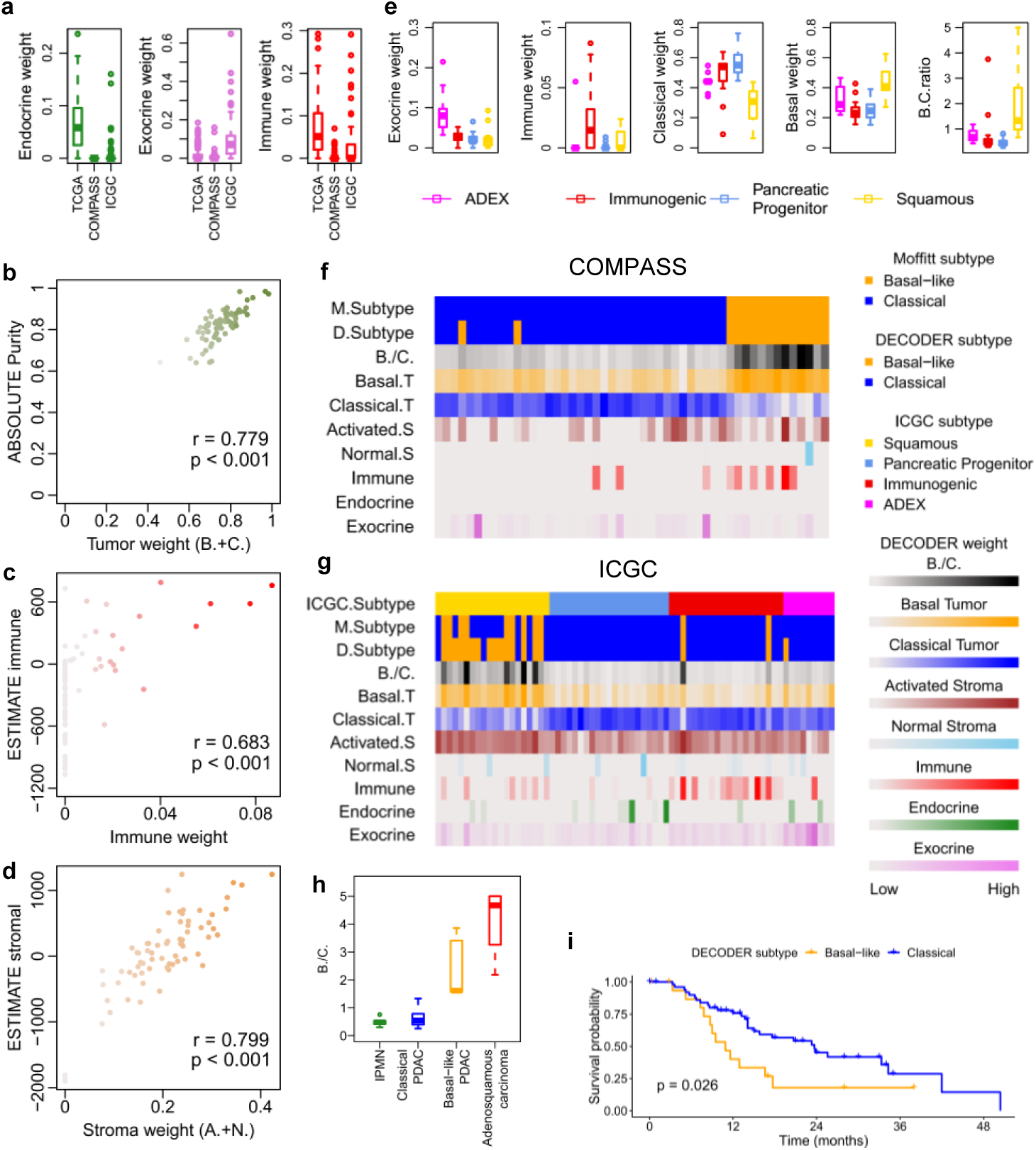
Estimated single sample compartment weights in COMPASS and ICGC pancreatic cancer samples. **a** Comparison of endocrine, exocrine and immune compartment weights in the COMPASS (microdissected), ICGC and TCGA PAAD datasets. **b, c &d** Correlations of DECODER estimated tumor weight (the sum of basal and classical tumor weights), immune weight and stroma weight (the sum of activated and normal stroma weights) with ABSOLUTE tumor purity, ESTIMATE immune score and ESTIMATE stromal score respectively. **e** Exocrine, immune, classical and basal compartment weights in ADEX, Immunogenic, Pancreatic Progenitor and Squamous subtypes in the ICGC dataset. **f & g** Estimated DECODER compartment weights, the ratio between basal and classical compartment weights (B./C.), tumor subtypes called by B./C. (D.Subtype) and tumor subtypes called by using Moffitt tumor genes (M.Subtype) are shown for COMPASS and ICGC samples. For COMPASS, samples were ordered by consensus clustering using Moffitt tumor genes. For ICGC, samples are ordered by ICGC subtypes. **h** B./C. for ICGC samples grouped by intraductal papillary mucinous neoplasm (IPMN), Classical PDAC, Basal-like PDAC and adenosquamous carcinoma. **i** Kaplan-Meier plot of DECODER tumor subtypes in the ICGC dataset.

In both the COMPASS and ICGC datasets, the basal weight, classical weight, as well as the ratio between them (*B./C.*) are associated with the consensus clustering based subtypes (Fig. 3f,g, Supplementary Fig. S3c-f). As expected, invasive intraductal papillary mucinous neoplasm (IPMN) shows the lowest *B./C.* while the adenosquamous carcinoma samples show the highest, suggesting that the *B./C.* ratio is able to identify the extremes of tumor histology^23^ (Fig. 3h). In addition, the DECODER derived basal and classical weights are differentially associated with patient outcome in ICGC, where survival data is available (Fig. 3i). These results demonstrated that the DECODER sample weights may faithfully and accurately recapitulate the differences in the biological make-up of a single sample.

### Application on TCGA PanCan ATAC-seq dataset

Previous PanCan analysis on an ATAC-seq dataset of 23 human cancers has identified 18 distinct clusters of samples, which showed strong concordance with the published multiomic iCluster scheme^24,25^. These studies found both homogeneous clusters for single tumor types, and heterogeneous clusters formed by mixed tumor types arising from the same organ systems or with similar features, with some cancer types split into multiple clusters. We therefore interrogated whether DECODER can deconvolve the PanCan ATAC-seq dataset containing 759 replicates from 410 unique samples into compartments that reflect inner biological composition. DECODER identified 22 primary compartments, the sample weights for which were subjected to unsupervised hierarchical clustering revealing overall alignment for 19 of them with cancer types, ATAC-seq-based clusters and iClusters (Fig. 4a). We then annotated the compartments by examining the associations of sample weights with cancer types, organ systems and previous cluster-base calls, and successfully associated 19 of them (Fig. 4b-d).

**Fig. 4.**
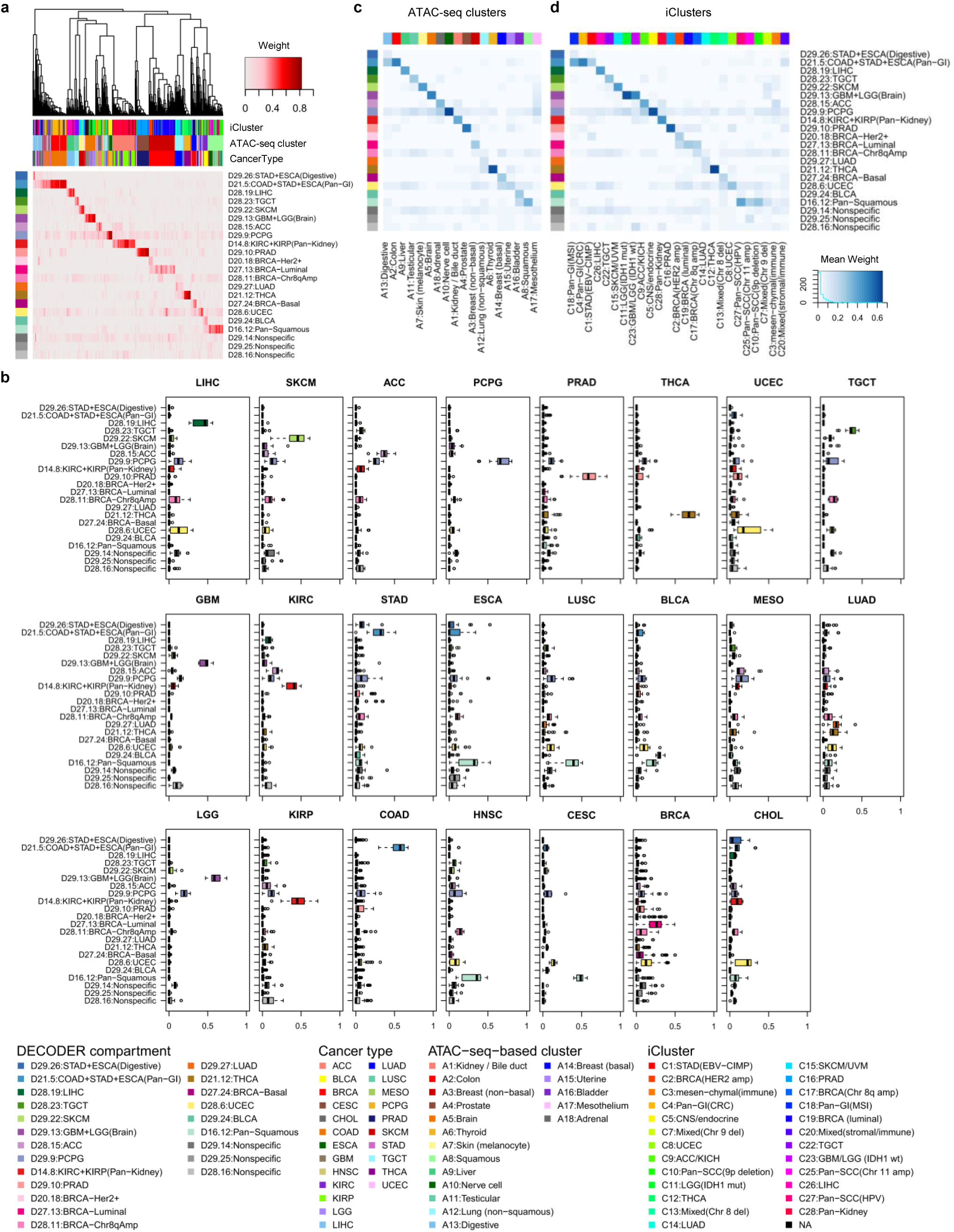
De novo deconvolution on TCGA ATAC-seq PanCan dataset. **a** Unsupervised hierarchical clustering on samples (columns on the heat map) using compartment weights. Cancer types, ATAC-seq-based cluster calls and iCluster calls for the samples are shown as tracks. Compartments (rows on the heat map) were manually ordered so that the enriched of weights are shown on the diagonal. The same order of the compartments is maintained in **b-d**. **b** Compartment weights for each cancer type. **c & d** The means of DECODER compartment weights relative to ATAC-seq-based clusters and iClusters.

Ten compartments were found to show uniquely higher weights in single cancer types, namely liver hepatocellular carcinoma (D28.19:LIHC), skin cutaneous melanoma (D29.22:SKCM), adrenocortical carcinoma (D28.15:ACC), pheochromocytoma and paraganglioma (D29.9:PCPG), prostate adenocarcinoma (D29.10:PRAD), thyroid carcinoma (D21.12:THCA), uterine corpus endometrial carcinoma (D28.6:UCEC), testicular germ cell tumors (D28.23:TGCT), lung adenocarcinoma (D29.27:LUAD) and bladder urothelial carcinoma (D29.24:BLCA) (Fig. 3b). Similarly, we found the sample weights for these compartments to be dominantly highest for ten respective ATAC-seq clusters and iClusters (Fig. 4c,d). This suggests that DECODER identified these compartments to be specifically associated with ten cancers, similar to the conclusion in ATAC-seq-based clustering and iCluster where these tumors dominated separate clusters.

DECODER also identified organ system associated compartments. For instance, brain (D29.13), pan-kidney (D14.8), pan-gastrointestinal (pan-GI) (D21.5), digestive (D29.26) and pan-squamous (D16.12). For both of the brain tumors (glioblastoma multiforme [GBM] and Brain Lower Grade Glioma [LGG]), D29.13 show exclusively high weight. Intriguingly, the compartment for PCPG (D29.9) also show higher weight than the rest for GBM and LGG, reflecting the fact that they are more anatomically similar. Comparing to ATAC-seq clusters and iClusters, D29.13 is associated with A5:Brain, and C11:LGG(IDH1 mut) and C23:GBM/LGG(IDH1 wt) respectively. D14.8 is distinctly highly weighted for kidney clear cell carcinoma (KIRC) and kidney renal papillary cell carcinoma (KIRP), represented as a pan-kidney compartment, which is associated with A1: Kidney/Bile duct and C28:Pan-Kidney. Similarly, D21.5:COAD+STAD+ESCA(Pan-GI) and D29.26:STAD+ESCA(Digestive) were found to be related to the pan-GI system and clusters, with D21.5 represented as showing the highest weights in colon adenocarcinoma (COAD) and stomach adenocarcinoma (STAD), and the second highest weights in esophageal carcinoma (ESCA). For ESCA, which may have squamous morphology components, the compartment with highest weight was annotated as the pan-squamous compartment (D16.12), showing the strongest associations with the squamous clusters in ATAC-seq and iCluster as well. The pan-squamous compartment (D16.12) is also the most highly represented in head and neck squamous cell carcinoma (HNSC), lung squamous cell carcinoma (LUSC) and cervical and endocervical cancers (CESC), known to have squamous histologies. Interestingly, while the squamous compartment shows the second highest weights in BLCA, we found the cancer specific D29.24:BLCA to be overrepresented as well. This is in agreemnet with the finding that BLCA has very diverse iCluster memberships^25^.

Compartment D27.24:BRCA-Basal, D20.18:BRCA-Her2+, D28.11:BRCA-Chr8qAmp and D27.13:BRCA-Luminal were all found be associated with breast invasive carcinoma (BRCA) (N=141), enabled by sample sufficiency in the dataset as expected by its known heterogeneity^26^. No specifically associated compartments were found in cholangiocarcinoma (CHOL, N=10) and mesothelioma (MESO, N=13) which may be confounded by the small samples sizes available. Similar for TGCT (N=18), although D28.23:TGCT was annotated to be specifically associated with it, we found other compartments showing noticeably high weights as well, which may due to the ineffective factorization of the compartment caused by insufficient samples.

## Discussion

DECODER is a novel and integrated framework, which we developed to perform *de novo* compartment deconvolution for any dataset with non-negative values, and conduct efficient compartment weight estimation for even a single sample of cancer types in TCGA. Standard NMF methods pre-define the number of factors (*K*) and assume the presence of *K* compartments in a dataset. In contrast, the *de novo* compartment deconvolution of DECODER is fluid and allows each of the compartments in a dataset to be identified at runs of different *K*, allowing the identification of each compartment to be more robust. This obviates the need for prior knowledge of the compartments and the number of them in a dataset, since compartments vary in different cancers or tissues and certain compartments present in one may be absent in another.

We applied DECODER to deconvolve each of the 33 cancer types in the TCGA RNA-seq datasets. Our results will facilitate the acquisition of cancer type specific marker genes. This may lead to more accurate estimation of certain compartment fractions, since as far as we know, current methods for deconvolution of tumor, stroma and immune fractions use the same set of curated marker genes for all cancer types^7,14^. Furthermore, cancer-specific marker genes may be studied as cancer specific biomarkers. The deconvolved mRNA compartments and respective marker genes for 33 cancer types are readily accessible online.

Our detailed examination on the resultant compartments of TCGA PAAD dataset demonstrated the robust identification of biological compartments in PDAC, as defined by previously known knowledge. In addition to the previously described compartments of Basal-like tumor, Classical tumor, Activated stroma, Normal stroma, immune, endocrine and exocrine, we also identified two new compartments, which we annotated as olfactory and histone. Interestingly, a study has associated the olfactory transduction pathway with pancreatic cancer risk^27^, and additional studies have shown the overexpression of olfactory receptors in multiple cancer types, including prostate^28,29^, bladder^30^ and breast^31^ cancers. In agreement with these studies, we found that our olfactory compartment was indeed present in prostate adenocarcinoma (PRAD), bladder urothelial carcinoma (BLCA), as well as several other cancer types (Supplementary Data 1). Similarly, the histone compartment was identified in multiple cancer types. Further investigation will be required to determine if the olfactory and histone compartments are true biological compartments or artifacts of inherent noise in RNA sequencing.

Unlike regular clustering-based methods, DECODER provides the possibility of examining a sample multidimensionally via the compartment weights, instead of forcing a given sample to a specific cluster. This provides more detailed information regarding the biological composition of a sample, which can be used for comparison across samples and datasets. In PDAC, absolute consensus for transcriptomic subtypes by different taxonomies has not been achieved, with the Exocrine-like/ADEX and Immunogenic subtypes at the center of the controversies^23^. By applying DECODER to the COMPASS and ICGC datasets, we demonstrated that in COMPASS, where samples were microdissected, there is comparatively less of the exocrine and immune compartments. In ICGC, the exocrine compartment weights correlate with the Exocrine-like/ADEX subtype while immune compartment weights correlate with the Immunogenic subtype. These findings agree with the association of low purity samples with Exocrine-like/ADEX and Immunogenic subtypes^19^ and suggest that the Exocrine-like/ADEX and Immunogenic subtypes may be explained by the presence of the non-tumor compartments. In addition, in the more heterogenous ICGC dataset, DECODER weights of basal and classical compartments correlate well with IPMN vs adenoquamous carcinomas. Therefore, DECODER may facilitate the better elucidation of the underlying biology of molecular subtypes.

Furthermore, relying on the results from the initial *de novo* deconvolution in TCGA PAAD, compartment weight estimation in ICGC and COMPASS was single-sample based, and therefore feasible in the clinical setting. Similarly, for any of the 33 cancer types in TCGA, DECODER can be used to estimate the compartment weights for a new given sample (i.e., sample weights) without the need to perform *de novo* deconvolution. Thus, DECODER is a powerful tool to break down a new tumor sample with known origin.

DECODER may be applied to any solid or hematologic malignancy, as well as liquid biopsies, and may also be used for platforms other than gene expression. From a computational perspective, DECODER could be applied to any data type with non-negative values derived from multiple platforms, e.g. ChIP-seq data, copy number variations, DNA methylation data and single cell sequencing platforms. As a proof of concept, we applied DECODER on the PanCan ATAC-seq dataset containing 23 cancer types in a combined fashion, and identified compartments associated with cell-of-origin and organ systems, which highly reproduced previous clustering-based interpretations of the PanCan analysis^24,25^. Similar to our application of DECODER in RNA-seq, deconvolution of the PanCan ATAC-seq dataset can also be applied to new samples. One limitation of our PanCan analysis may be the unequal number of samples in different cancer types (range of number of samples: 7 to 141), which may lead to an unbalanced number of compartments identified for different cancer types. Therefore, it is possible that more unbiased compartment information may be derived in the setting of a greater number of samples especially for the cancers with smaller sample sizes.

From a clinical perspective, the ability of DECODER to identify sample compartments without the *a priori* assumption of an actual number of compartments, makes it a powerful tool for comparing tumor heterogeneity across samples and for the very challenging clinical conundrum of cancers of unknown primary. Cancers of unknown primary are metastatic cancers where the primary site of origin cannot be identified. Optimal treatment of these cancers remains a challenge as knowledge of the primary site guides treatment decisions. With carefully curated inclusion of combined samples for *de novo* deconvolution, application of DECODER across multiple cancer types using data from various platforms may provide more biological information that can better guide treatment for cancers of unknown primary.

## Methods

### Data preparation

A published collection of microarray data (GSE71729, GSE21501) from a previous study^16^ was used for comparison between *de novo* deconvolution of DECODER and previous NMF runs in the study. To build a resource base of deconvolved compartment information for cancers, TCGA normalized RNA-seq gene expression data from 33 cancer types were downloaded from the Broad Institute FIREHOSE portal (http://gdac.broadinstitute.org). For the PAAD dataset, out of 183 samples deposited in the portal, we included 150 PDAC samples for further analysis^19^. For the rest of the cancer types, all of the downloaded samples were used for the *de novo* deconvolution analysis. The normalized ATAC-seq counts within the pan-cancer peak set were downloaded (https://gdc.cancer.gov/about-data/publications/ATACseq-AWG). Loci with mean counts across samples greater than the overall mean, and standard deviation (SD) of counts across samples greater than the mean of all SD (N=105,138) were subjected to *de novo* deconvolution of DECODER.

For single sample compartment weight estimation analysis, 70 pancreatic cancer samples with matched mRNA subtype calls from Bailey et al.^18^, and RNA-seq data, as well as clinical data (PACA-AU) were downloaded from ICGC data portal (http://dcc.icgc.org/). Similarly, RNA-seq data of 50 PDAC samples that underwent laser capture microdissection in COMPASS trial were involved in this analysis^20^.

A summary for datasets involved in this study is provided in Supplementary Table S1.

### NMF seed training

To handle the stochastic nature of the NMF, a stable gene weight seed was trained before the application of a final NMF. For *de novo* deconvolution of microarray and RNA-seq expression profiles, highly expressed genes in the top third quartile were subjected to selection of the 5,000 most variable ones. Then 80% random samples were used at each run of 10,000 iterations, in which top genes are derived for each factor. Finally, the frequency of the genes to be determined as the top genes for each factor was summarized as the seed matrix W’’. Using this gene weights seed W’’, a final NMF was applied to the whole dataset, with a robust deconvolution of gene weights (W’) and samples weights (H’). Similarly, 8,000 most highly expressed and variable ATAC-seq peaks were involved in the seed training process.

### Sample weights and gene weights projection

Given the gene weight matrix (W’) and a column/sample (y) of expression matrix (A), non-negative least square (NNLS) was used to solve the problem of finding a vector (x) that minimized the Euclidean norm of Ax-y. x represents the compartment weights for each sample, resulting in the identification of sample weight matrix (H). Similarly, NNLS was then used to project H onto each of the genes resulting in gene weights (W).

In single sample weight estimation, NNLS algorithm was used on the deconvolved gene weights (W) and each column/sample in a given expression matrix (B) to identify compartment weights for each single sample. For pancreatic cancer data in both *de novo* deconvolution and single sample weight estimation, the compartment weights of each sample for basal tumor, classical tumor, activated stroma, normal stroma, immune, endocrine and exocrine are normalized so that the sum of them equals to one.

### Gene analysis and compartment annotation

For a factor (*i*th column in W) at each run of *K*, the genes were ranked in descending order of the weight difference between the loading value in the *i*th column and the largest loading value in the rest of the columns. For each compartment, the marker genes were identified as the top genes with normalized weight differences exponentially greater than the rest, which are above the geometric inflection point on the ranking curve (Supplementary Fig. 1b). Ranked genes in each compartment were subjected to gene set analysis using annotated gene sets from MSigDB v3.1^21^, assessed by the Kolmogorov-Smirnov statistic with Benjamini-Hochberg correction for significance as described before^16^. For RNA-seq data, final compartments of TCGA PAAD are annotated based on MSigDB terms and prior knowledge; and for the rest of the cancer types, the immune, stroma, basal, olfactory and histone compartments are annotated according to MSigDB as well, with the rest of the compartments unannotated.

### Factor score calculation and linkage establishment

The overlapping percentages of top 250 genes between each of the factors at *K*=*k* and each of the factors at *K*=*k*-1 were calculated. For the *i*th factor at *K*=*k*, if the *ii*th factor at *K*=*k*-1 showed the largest overlapping and it was greater than 0.1, then the overlapping percentage was determined as the score for the *i*th factor at *K*=*k*. And these two factors were considered as the same compartments at adjacent runs of *K*, thus a link was established between them. However, if the largest overlapping percentage between the *i*th factor at *K*=*k* and the factors at *K*=*k*-1 was smaller than 0.1, the *i*th factor at *K*=*k* was considered a newly emerged factor at *K*=*k*, not showing up at *K*=*k*-1 yet. Similarly, for the *i*th factor at *K*=*k*, if no linkage was established between this factor and any of the factors at *K*=*k*+1, the respective factor was considered too split or diluted to be detected in the resolution of *K*=*k*+1.

### Compartments identification from factors

The linking of all the similar factors at multiple runs of *K* leads to the placement of all the identified factors in a dataset on a tree-like plot. To determine the optimal factor along each link, the pattern of the factor scores was considered, so that a resultant compartment a) is on a link with more than 2 factors on it, b) shows a score greater than the third quartile (Q^3^) of all the scores, c) has more than two continuous (adjacently linked) (primary) or one (unstable) factor(s) greater than the median quartile (Q^2^) of all the scores, d) is with the maximum score among the continuous factors with score higher than Q^2^ (candidates in a high-score factor block) and e) has the largest number of factors in the high-score block or has the greatest score if multiple high-score blocks and candidates found. Finally, if a primary compartment is found to have over 100 overlaps in top 250 genes with another primary compartment, and both of them are on the same link, the compartment identified at greater *K* is then labeled as a ‘secondary’ compartment.

### Ten-fold cross validation

In TCGA PAAD, samples were randomly partitioned into ten subsets. *De novo* deconvolution was then performed on each combination of nine-fold of samples to derive final gene weights. Then for each one-fold of samples, the sample weights were then estimated by the gene weights derived by the other nine-fold of samples by the non-negative least square (NNLS) algorithm. Finally, the estimated sample weights were compared to those derived from performing *de novo* deconvolution on the whole dataset.

### Statistical information

Paired Wilcoxon rank-sum test was used for comparison of basal and classical weights in subtyped samples. Wilcoxon rank-sum test was used to compare ratios and differences of basal and classical weights in subtyped samples. In correlation analysis, Spearman correlation was used to compare DECODER derived weights with ESTIMATE derived scores where the scales are largely different. Otherwise, Pearson correlation was involved. For survival analysis, continuous variables were analyzed by Cox proportional hazards regression model and categorical variables were analyzed by log-rank test.

### Usage of DECODER and code availability

The configure files for *de novo* deconvolution (Moffitt microarray, TCGA RNA-seq and TCGA PanCan ATAC-seq datasets), and single sample weight estimation (COMPASS PDAC and ICGC pancreatic cancer datasets) are provided as Supplementary Table S2-S6.

A Matlab repository for DECODER is available online, with the utilities of *de novo* deconvolution and single sample compartment weight estimation. All the downloaded datasets are also deposited in this repository, along with all the deconvolution results in this study. In addition, an R package for single sample compartment weight estimation for TCGA cancers is also readily accessible online.

## Supporting information

Supplementary Data 1 Combined_compartment_info

Supplementary Data 2 TCGA_PAAD_marker_genes

Supplementary Data 3 Sample_weights

