## Supplementary Information for "*<u>De</u> novo* <u>co</u>mpartment <u>de</u>convolution and weight estimation of tumo<u>r</u> samples (DECODER)"

### ***De novo* compartment deconvolution and weight estimation of tumor samples (DECODER)**

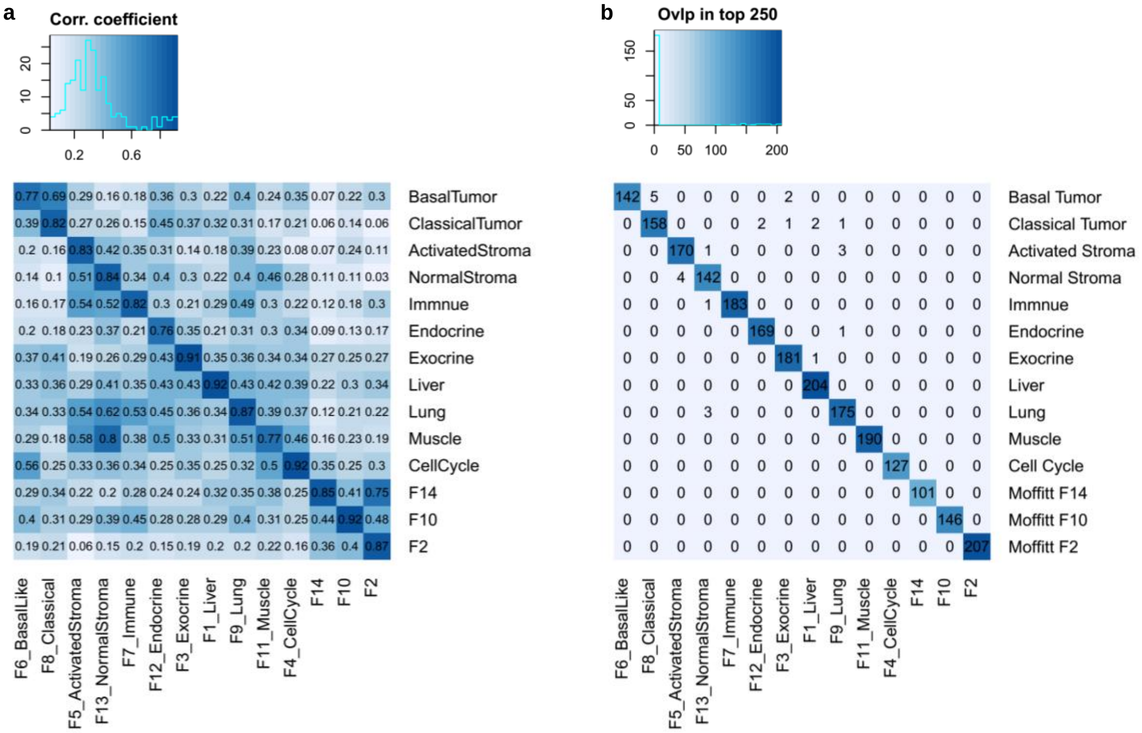


**Supplementary Figure S1**. Comparison between factors identified in the previous study using empirical number of factors (***K***=14) and primary compartments identified by the *de novo* deconvolution of DECODER in the Moffitt microarray dataset. **a** Correlation between gene weights. **b** Number of overlaps in top 250 genes.

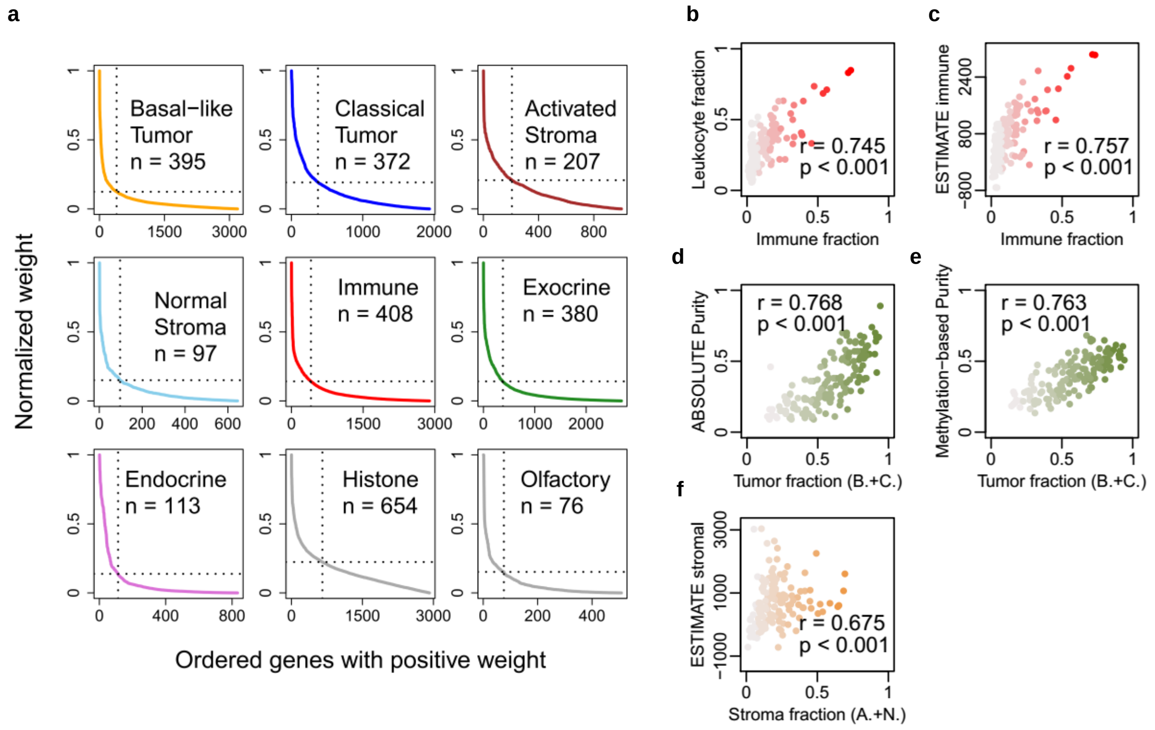


**Supplementary Figure S2**. Marker genes of compartments in TCGA PAAD. **a** Identification of marker genes for each compartment. **b & c** Correlation between the immune fraction calculated using marker genes for the immune compartment, and the leukocyte fraction and ESTIMATE immune score. **d & e** Correlation between the tumor fraction (sum of basal and classical fraction) and the tumor fraction estimated by ABSOLUTE and methylation. **f** Correlation between the stroma fraction (sum of activated and normal fraction) and the ESTIMATE stromal score.

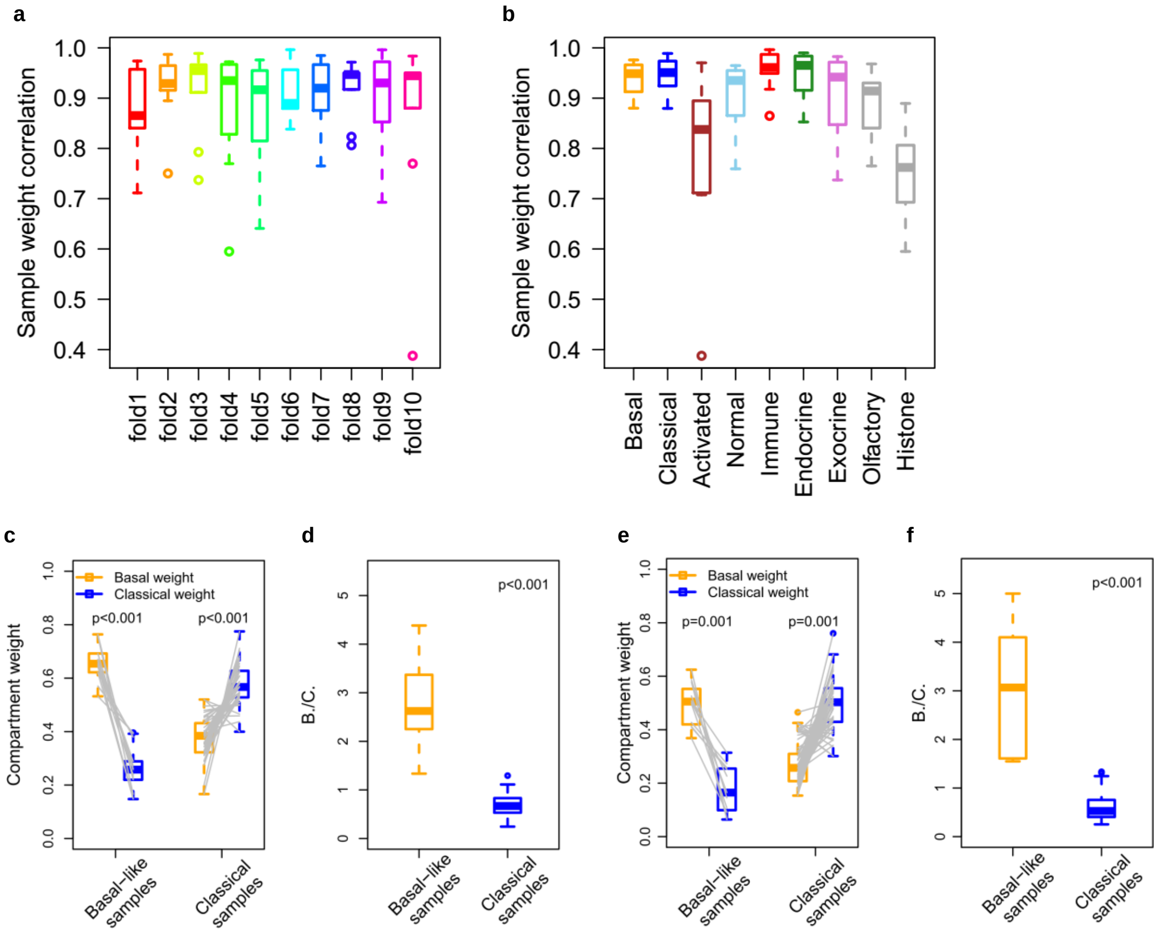


**Supplementary Figure S3**. Single sample weight estimation. **a & b** Ten-fold cross validation comparing sample weights derived from *de novo* deconvolution and sample weights estimated using trained gene weights by the non-negative least square (NNLS) algorithm. **c & e** Basal weight, classical weight and the ratio between them (B./C.) in Basal-like and Classical samples subtyped by previous consensus clustering based method in the COMPASS dataset. **d & f** Basal weight, classical weight and the ratio between them (B./C.) in Basal-like and Classical samples subtyped by previous consensus clustering based method in the ICGC dataset.

**Supplementary Table S1**. **Datasets involved in this study.**

| Dataset | Description | Link |
|---|---|---|
| Moffitt Microarray | Primary tumor, metastatic and normal samples | https://www.ncbi.nlm.nih.gov/geo/query/acc.cgi?acc=GSE71729 https://www.ncbi.nlm.nih.gov/geo/query/acc.cgi?acc=GSE21501 |
| TCGA RNA-seq | 33 cancer types | http://gdac.broadinstitute.org |
| COMPASS PDAC | Microdissected PDAC | https://www.ebi.ac.uk/ega/studies/EGAS00001002543 |
| ICGC Pancreatic Cancer | Panceatic cancers including PDAC, IPMN, adenoquamous carcinomas and acinar cell carcinomas | http://dcc.icgc.org/ |
| TCGA ATAC-seq | 23 cancer types | https://gdc.cancer.gov/about-data/publications/ATACseq-AWG |

**Supplementary Table SS**. **Configure file for de novo deconvolution of the Moffitt microarray dataset.**

| | |
|---|---|
| dataType | Microarray |
| dataMatrix | Moffitt_PDAC_array.mat |
| dataFormat | mat |
| geneIDType | geneSymbol |
| logTransformed | yes |
| rangeK | 2:25 |
| repTimes | 10000 |

**Supplementary Table S3**. **Configure file for de novo deconvolution of the TCGA RNA-seq datasets.**
TCGA pancreatic adenocarcinoma (PAAD) RNA-seq data is provided as demo_data.tsv.

| | |
|---|---|
| dataType | RNAseq |
| dataMatrix | demo_data.tsv |
| dataFormat | tsv |
| geneIDType | TCGA |
| logTransformed | no |
| rangeK | auto |
| repTimes | 10000 |

**Supplementary Table S4**. **Configure file for de novo deconvolution of the TCGA ATAC-seq PanCan dataset.**

| | |
|---|---|
| dataType | ATACseq |
| dataMatrix | TCGA_ATACseq.mat |
| dataFormat | mat |
| geneIDType | |
| logTransformed | yes |
| rangeK | 2:30 |
| repTimes | 10000 |

**Supplementary Table S5**. **Configure file for single sample weight estimation of the COMPASS dataset.**

| | |
|---|---|
| refSet | TCGA_RNAseq_PAAD |
| dataMatrix | COMPASS_PDAC.tsv |
| dataFormat | tsv |
| geneIDType | geneSymbol |
| logTransformed | no |

**Supplementary Table S6**. **Configure file for single sample weight estimation of the ICGC dataset.**

| | |
|---|---|
| refSet | TCGA_RNAseq_PAAD |
| dataMatrix | ICGC_PancreaticCancer.tsv |
| dataFormat | tsv |
| geneIDType | geneSymbol |
| logTransformed | no |
